# Evolving small-molecule biosensors with improved performance and reprogrammed ligand specificity using OrthoRep

**DOI:** 10.1101/2021.06.15.448565

**Authors:** Alex A. Javanpour, Chang C. Liu

**Affiliations:** Department of Biomedical Engineering, University of California, Irvine, CA 92697, USA; Center for Synthetic Biology, University of California, Irvine, CA 92697, USA; Department of Chemistry, University of California, Irvine, CA 92697, USA; Department of Molecular Biology & Biochemistry, University of California, Irvine, CA 92697, USA

## Abstract

Genetically-encoded biosensors are valuable for the optimization of small-molecule biosynthesis pathways, because they transduce the production of small-molecule ligands into a readout compatible with high-throughput screening or selection *in vivo*. However, engineering biosensors with appropriate response functions and ligand specificities remains challenging. Here, we show that the continuous hypermutation system, OrthoRep, can be effectively applied to evolve biosensors with high dynamic range, reprogrammed activity towards desired non-cognate ligands, and proper operational range for coupling to biosynthetic pathways. In particular, we encoded the allosteric transcriptional factor, BenM, on OrthoRep such that propagation of host yeast cells resulted in BenM’s rapid and continuous diversification. When these cells were subjected to cycles of culturing and sorting on BenM activity in the presence and absence of its cognate ligand, muconic acid, or the non-cognate ligand, adipic acid, we obtained multiple BenM variants that respond to their corresponding ligands. These biosensors outperform previously-engineered BenM-based biosensors by achieving substantially greater dynamic range (up to ~180-fold-induction) and broadened operational range. Expression of select BenM variants in the presence of a muconic acid biosynthetic pathway demonstrated sensitive biosensor activation without saturating response, which should enable pathway and host engineering for higher production of muconic and adipic acids. Given the streamlined manner in which high-performance and versatile biosensors were evolved using OrthoRep, this study provides a template for generating custom biosensors for metabolic pathway engineering and other biotechnology goals.

## Introduction

Microbial biosynthesis of small-molecules is an attractive alternative to traditional chemical synthesis for a wide array of therapeutically and commercially relevant compounds. For example, yeast have been utilized for the biosynthetic production of the anti-malarial precursor, artemisinic acid (1); a range of opioids, such as thebaine and hydrocodone (2); perfume agents, such as santalene (3); biofuels (4); and oleochemicals (5). However, engineering microbes to efficiently produce a target small-molecule often requires substantial modification of heterologous enzymes and host cell metabolism, necessitating search through a vast space of designs. Although large libraries of pathway and host modifications can be readily constructed using modern genetic engineering techniques, finding the most efficient producers is typically low throughput: agar plate screens on colonies, utilizing a visual output or growth complementation, are limited to small library sizes of up to 10^5^; and microtiter plate-based screening approaches coupled analytical techniques such as mass spectrometry and liquid/gas chromatography are limited to libraries of up to 10^4^ (6, 7).

To convert the identification, engineering, and evolution of metabolic pathways into a high throughput exercise, synthetic biologists have focused on the generation of custom genetically-encoded biosensors that can specifically report on the production levels of desired molecules *in vivo* (8–13). Properly engineered, such biosensors transduce the amount of a target small-molecule into a biological output, such as the strength of reporter gene expression. These outputs can then be used for sorting, as in the case of fluorescence reporters, or selection, as in the case of a selectable gene, forming the crucial link between an arbitrary molecule’s production and the high-throughput enrichment of pathways and host modifications that efficiently produce the molecule. The generation of custom biosensors has therefore emerged as an important area of research within the field of metabolic engineering.

A powerful class of potential custom biosensors are allosteric transcription factors (aTFs), which have an effector-binding domain responsible for recognizing cognate ligands and a DNA binding domain that modulates target gene transcription when the cognate ligand is bound (14, 15). Because of their modular architecture and abundance, natural aTFs are attractive starting points for engineering custom biosensor properties, such as regulatory response, regulatory logic, and specificity for non-cognate ligands (11, 12). Recently, Jensen, Keasling, and colleagues demonstrated that the largest family of transcriptional regulators found in prokaryotes, the LysR-type transcriptional regulators (LTTRs), are both portable into workhorse bioproduction hosts such as *Saccharomyces cerevisiae* and highly engineerable for custom biosensor behaviors (16, 17). For example, they showed that the homotetrameric prokaryotic LTTR, BenM, can activate transcription of genes of interest in *S. cerevisiae* when BenM’s cognate ligand, *cis,cis*-muconic acid (CCM), is present. This was done by engineering a common yeast promoter, CYC1p, to contain BenM’s DNA operator sequence, *BenO*. This hybrid CYC1p-BenO promoter was then used to control GFP expression, allowing for a series of directed evolution campaigns that yielded new BenM variants through fluorescence activated cell sorting (FACS). In these efforts, the Jensen Lab generated BenM variants with increased dynamic and operational range for CCM, inverted regulatory logic, as well as reprogrammed specificity for adipic acid (AA). For example, the most prominent BenM mutants identified in Snoek *et al.* (17) displayed a 15-fold increase in output level, a 40-fold shift in operational range, and complete inversion-of-function from ligand-induced activation to repression when compared to the parental BenM biosensor. However, there is need for additional BenM biosensors with expanded dynamic range, operational range, and higher specificity given the diversity of metabolic engineering contexts in which BenMs may be applied.

In this study, we sought to evolve high-performance BenM-based biosensors using the *in vivo* continuous hypermutation system, OrthoRep. By encoding BenM on OrthoRep, straightforward cycles of culturing and FACS in the presence and absence of CCM or AA yielded BenM variants with desired behaviors. Because BenM continuously diversifies when encoded on OrthoRep, we were able to implement many cycles of evolutionary improvement in a straightforward manner, resulting in superior multi-mutation BenM variants with high dynamic range, broad operational range, and improved specificity compared to literature benchmarks. This contrasts with past biosensor directed evolution strategies that carried out cycles of FACS enrichment on a static library of BenMs (16, 17), restricting the improvements accessible. Combinatorial examination of mutations found in our evolved BenM variants showed that all mutations characterized improved functionality in at least one context, validating the utility of OrthoRep-driven evolution cycles. Finally, we expressed our evolved BenM variants in the presence of a CCM production pathway to show operational sensing for the goal of evolving CCM and AA metabolic pathways. We argue that the demonstrated effectiveness of OrthoRep-driven BenM evolution combined with the inherent scalability of OrthoRep-based evolution experiments will advance future biosensor engineering campaigns, as the scope of desired biosensors, both in specific molecules sensed and performance across different operational ranges, is vast.

## Results

### Directed evolution of BenM with OrthoRep

We sought to demonstrate OrthoRep-driven biosensor evolution by 1) increasing the ability of BenM to sense CCM and 2) reprogramming BenM to respond to AA. To establish the BenM evolution strain, we started with a CEN.PK113-5A *S. cerevisiae* strain previously engineered to express GFP under the control of the hybrid CYC1p-BenO promoter, which is activated by BenM in the presence of CCM (MeLS0138). We fully deleted the *URA3* and *TRP1* genes in this parent strain and inserted OrthoRep components by protoplast fusion and transformation following established pipelines (18–20). The resulting BenM evolution strain includes a recombinant orthogonal p1 plasmid that encodes WT *BenM* along with a selectable marker (*URA3*) and a nuclear (*CEN6/ARS4*) plasmid that encodes the error-prone orthogonal DNA polymerase, TP-DNAP1-4-2, along with a selectable marker (*TRP1*) (**Figure 1A**). This evolution strain uses the error-prone orthogonal DNAP to continuously replicate BenM at a high mutation rate of 10^−5^ substitutions per base (spb) (19).

**Figure 1.**
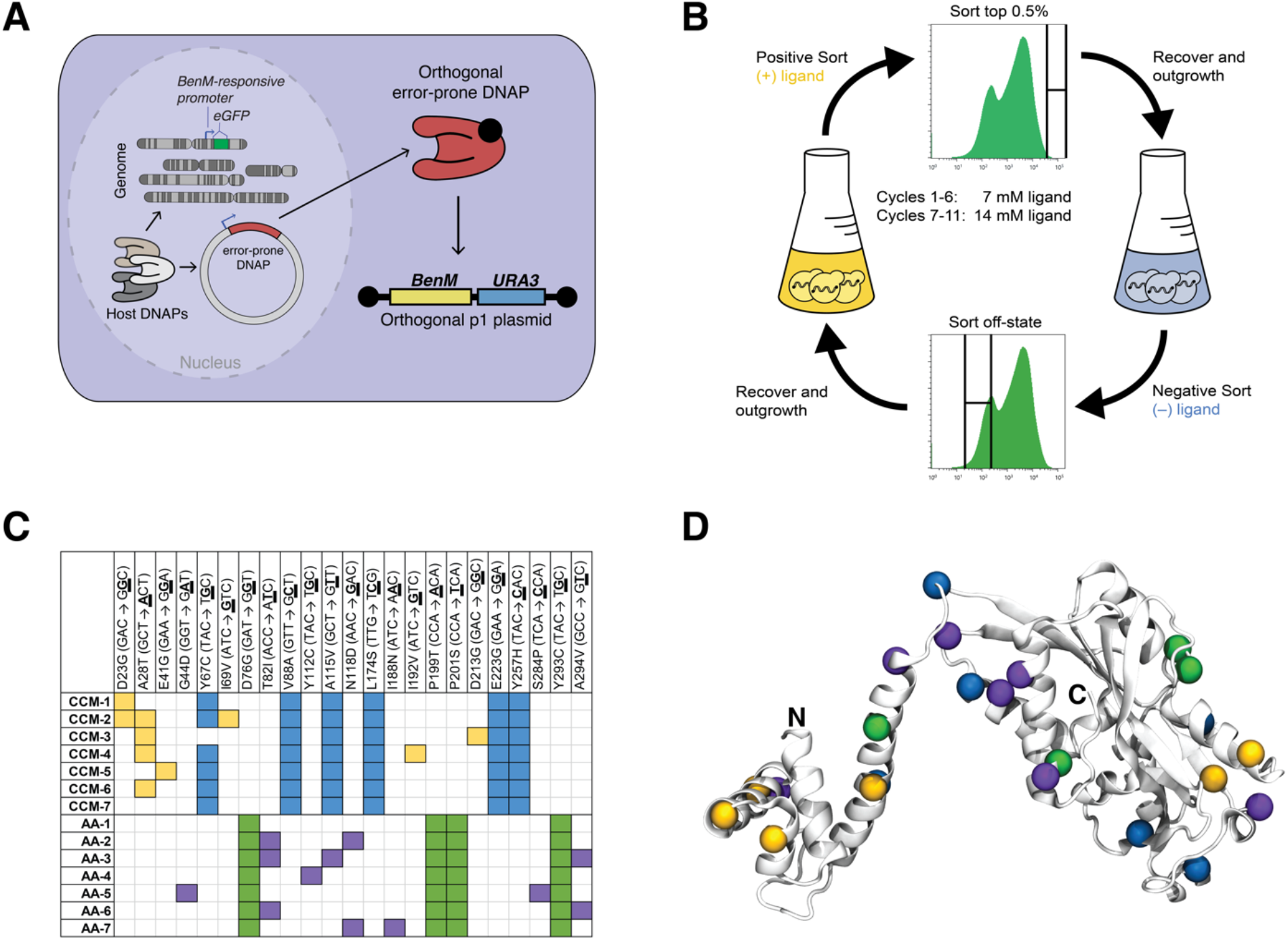
Evolution of BenM biosensors for *cis,cis*-muconic acid (CCM) and adipic acid (AA). (**A**) OrthoRep evolution strain. The accessory plasmid p2 is not shown for simplicity. (**B**) Overview of FACS-based evolution pipeline. (**C**) Tables showing mutations sequenced from top 7 CCM and AA hits. (**D**) Crystal structure of BenM (PDB: 3K1N) with mutations in (**C**) shown.

To evolve BenM, we first passaged the evolution strain for ~150 generations to accumulate BenM mutational diversity before introducing selection. Then, FACS-based evolution cycles were initiated. Each cycle consisted of 1) culturing cells in the presence of CCM or AA, 2) a positive sort where the top 0.5% most fluorescent cells were collected, 3) culturing cells in the absence of CCM or AA, and 4) a negative sort to remove any BenM mutants that constitutively activate GFP expression (**Figure 1B**). During culturing steps, BenM autonomously diversifies, allowing cycles of this process to improve BenM activity over time. Over the span of 11 total cycles, with the concentration of ligand increasing from ~7 mM to ~14 mM at cycle 7 to further broaden operational range, the overall behavior of each population adapted as desired: the average fluorescence of the population increased when cultured in the presence of CCM or AA and remained at background levels in the absence of CCM and AA. BenM alleles present in the cultures after cycle 11 were then isolated *via* PCR from population p1 DNA and integrated into the genome of a fresh yeast reporter strain as a single copy (*ura3Δ0::REV1p-BenM-tTDH1*) through transformation. The reporter strain, which does not include OrthoRep components, encodes GFP under the control of the hybrid CYC1p-BenO promoter. This allowed us to measure BenM activity in a context where BenM is encoded on the yeast genome, replicating how it would be applied as a biosensor for metabolic engineering and biotechnology goals. Ninety-six clones were randomly picked and tested for their ability to respond to ~14 mM CCM or AA, and the top 7 clones with highest dynamic range, defined as the fold-induction of GFP relative to the condition with no ligand present, were sequenced (**Figure 1C**).

The top 7 BenM variants evolved to sense CCM (CCM-1-7) contained between 6 and 9 non-synonymous mutations including a core set of five mutations shared among all variants (V88A, A115V, L174S, E223G, and Y257H) (**Figure 1C**). Additional mutations that some CCM biosensors contained include D213G, which is structurally located in the dimer-dimer interface of BenM (21, 22); and a near-consensus mutant, A28T, which was hypothesized in a prior structural study to enhance BenM’s interaction with the phosphate backbone of DNA (23). Specifically, it was suggested that replacing A28 with T or S could provide a hydroxyl to enhance BenM’s interaction with the phosphate backbone of DNA. Interestingly, other LTTRs contain a T or S at the position homologous to A28 (23). The top 7 BenM variants evolved to sense AA (AA-1-7) contained 5 to 8 non-synonymous mutations including a core set of four mutations shared among all variants (D76G, P199T, P201S, and Y293C) (**Figure 1C**). One of these mutations, P201S, is at a position involved in the dimer-dimer interface of BenM and was also uncovered in previous work evolving BenM to sense AA; it likely functions to change ligand specificity, as P201 forms an interaction with CCM (17, 22). Two other mutations, D76G and G44D, were notable because many LTTRs have a charged or polar amino acid at D76, and G44 is highly conserved (23). Although D76G and G44D could be neutral or deleterious hitchhiker mutations, we found this was not the case (see below). Overall, mutations were found throughout BenM, highlighting the wide range of interdomain mutations that can modulate allosteric regulation of BenM (**Figure 1D**).

### Evolved CCM and AA biosensors exhibit high dynamic range, operational range, and specificity

To fully characterize the performance of biosensors CCM-1-7 and AA-1-7, we obtained *in vivo* ligand response curves using the fluorescence of GFP driven by CYC1p-BenO as the readout (**Figures 2A** and **B**). Strains were grown in a range of ligand concentrations up to saturation in biological triplicate, and the dynamic range was determined for all concentrations (**Figures 2C** and **D**). For comparison, WT BenM and a set of benchmark BenM-based biosensors evolved in previous studies were tested alongside CCM-1-7 and AA-1-7. These benchmark biosensors include TM (H110R, F211V, Y286N) and MP17_D08 (A230V, F253S, Y286N, Y293H), which were previously evolved to sense CCM with improved dynamic range and expanded operational range, respectively (16, 17); and TiSNO120 (A130D, A153G, P201S, E287V), which was previously evolved for sensing AA (17).

**Figure 2.**
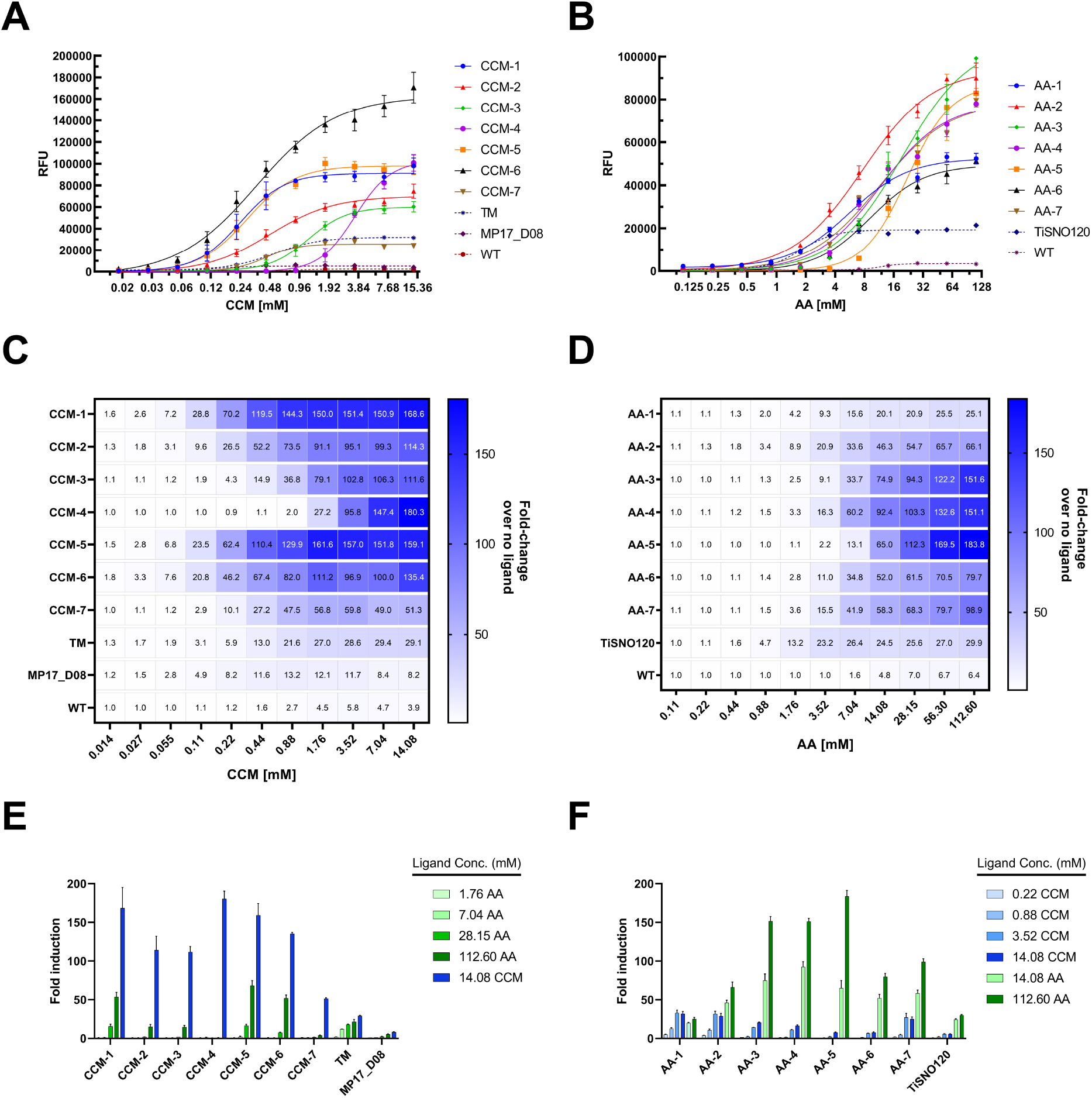
Performance of evolved CCM and AA biosensors. (**A**) CCM response curves for top 7 CCM biosensors. (**B**) AA response curves for top 7 AA biosensors. (**C** and **D**) Heat maps for CCM and AA biosensor response displaying mean fold-induction over the uninduced control (0 mM ligand) for CCM (**C**) and AA (**D**). (**E** and **F**) CCM and AA biosensor responses for off-target ligand. For all panels, data were collected in biological triplicate, and the means ± one standard deviation (error bar) are shown.

Nearly all variants evolved in this study outperformed benchmark biosensors in their dynamic range and have comparable or broadened operational ranges that span multiple orders of magnitude (**Figures 2A** and **B**). For example, CCM-4 had a dynamic range of ~180-fold and CCM-6 had a dynamic range of ~135-fold with a graded response over at least ~0.02-14 mM CCM. This contrasts with TM, with a dynamic range of ~29-fold and a response that saturates at ~2 mM CCM, and MP17_D08, with a dynamic range of ~13-fold and a response that saturates at ~1 mM CCM. Likewise, AA-5 had a dynamic range of ~180-fold and AA-3 had a dynamic range of ~150-fold with a graded response over at least ~1.5-112 mM AA. This contrasts with TiSNO120 with a dynamic range of ~30-fold and a response that saturates at ~3.5 mM AA. Notably, the large operational range of our AA biosensors, with several capable of discriminating up to ~110 mM AA, resulted from evolution campaigns that only used up to ~14 mM AA. This suggests that continued evolution of our AA biosensors may yield variants that are sensitive to even lower concentrations of AA, although these would have limited value for metabolic pathway evolution, because existing production pathways can already yield AA above the minimum concentrations our AA biosensors can detect (24).

Several BenM variants we evolved also exhibited high specificity for their cognate ligands compared to benchmark biosensors. For example, CCM-4 had no detectable activity for AA, in contrast to TM and MP17_D08 (**Figure 2E**). Likewise, AA-5 and AA-6 responded poorly to the non-cognate ligand, CCM, such that the saturation response for AA was ~24- and ~10-fold higher than the saturation response for CCM, respectively. This contrasts with TiSNO120 where the saturation response for AA is ~5-fold higher than the saturation response for CCM (**Figure 2F**). Notably, we did not perform any negative selection against non-cognate ligands in our evolution campaigns. Therefore, observed improvements in specificity were likely the byproduct of the many cycles of functional evolution enabled by OrthoRep, which may have resulted in specialization and associated specificity for the cognate ligand. Taken together, our OrthoRep-evolved BenMs form a new set of *in vivo* biosensors for CCM and AA distinguished by their diverse and exceptional performance. This should translate to versatility in their future application.

### Analysis of biosensor mutations reveals complex contributions to function

We next sought to determine how individual and collections of mutations in our evolved CCM and AA biosensors contribute to their functions. Because the evolved CCM-1-7 and AA-1-7 BenM variants each contain 4-9 mutations, it was impractical to analyze all possible mutant combinations. Instead, we divided mutations into two categories: those shared by (almost) all alleles (consensus) and those that were unique to only some alleles (secondary). For CCM-1-7, 5 mutations were shared among all variants and 6 mutations were shared among 6 of the 7 variants, so we chose to include 6 mutations as the consensus set: Y67C, V88A, A115V, L174S, E223G, and Y257H or CAVSGH for short. Likewise for AA-1-7, 4 mutations were shared among all variants, defining the consensus set: D76G, P199T, P201S, and Y293C or GTSC for short. We then carried out a series of studies focusing primarily on the CAVSGH and GTSC consensus sequences and, in select cases, the combination of consensus with secondary mutations. Yeast reporter strains with GFP driven by CYC1p-BenO were engineered to encode genomically-integrated biosensor variants (*ura3Δ0::REV1p-BenM_Variant-tTDH1*) containing the various mutant combinations of interest and GFP fluorescence was measured for all strains in the presence of no, medium, or high levels of CCM (0 mM, 0.88 mM or 14.08 mM) and AA (0 mM, 7.04 mM, or 112.60 mM) (**Figure 3**).

**Figure 3.**
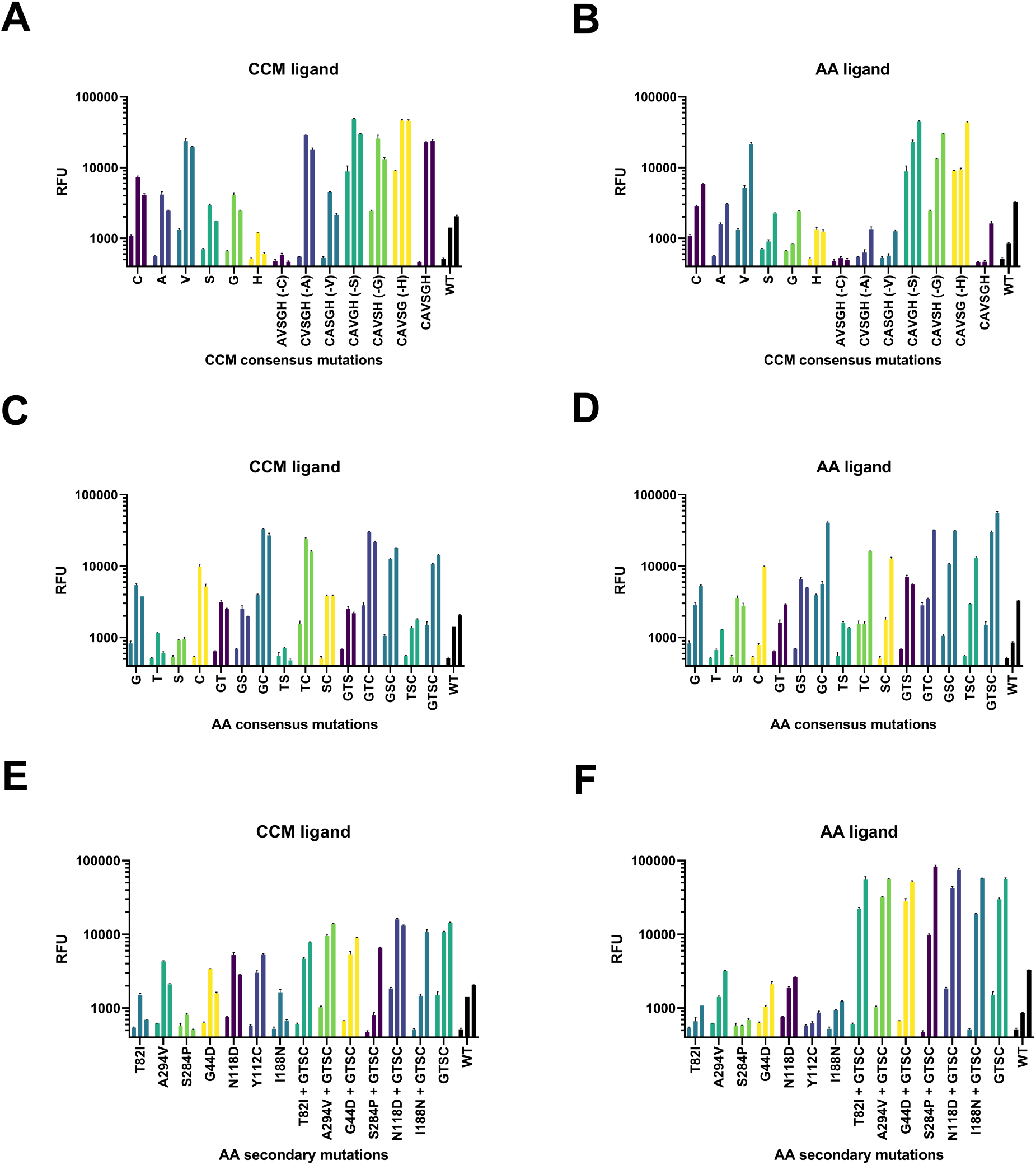
Combinatorial analysis of mutations observed in top 7 CCM and AA hits. (**A**) All 6 CCM consensus mutations as single or quintuple mutants induced with 0 mM (first bar in each set), 0.88 mM (second bar in each set), or 14.08 mM (third bar in each set) CCM. The consensus mutation that was removed to yield a given quintuple mutant is listed in parenthesis. (**B**) Same as (**A**), but induced with 0 mM, 7.04 mM, or 112.60 mM AA. (**C**) All combinations of the 4 AA consensus mutants induced with 0 mM (first bar in each set), 0.88 mM (second bar in each set), or 14.08 mM (third bar in each set) CCM. (**D**) Same as (**C**), but induced with 0 mM, 7.04 mM, or 112.60 mM AA. (**E**) Secondary AA mutants induced with 0 mM (first bar in each set), 0.88 mM (second bar in each set), or 14.08 mM (third bar in each set) CCM. (**F**) Same as (**E**), but induced with 0 mM, 7.04 mM, or 112.60 mM AA. Consensus mutation abbreviations for (**A**) and (**B**): C, Y67C; A, V88A; V, A115V; S, L174S; G, E223G; H, Y257H. Consensus mutation abbreviations for (**C**)-(**F**): G, D76G; T, P199T; S, P201S; C, Y293C. Data were collected in biological duplicate, and the means ± range (error bar) are shown.

For CCM biosensors, we tested how each mutation in the consensus set contributes to function as an individual mutation in BenM or if removed from the consensus. We found that a set of mutations (C, A, V) increased biosensor response in the presence of CCM but did so in a way that also increased background activity in the absence of ligand as well as activity in the presence of the non-cognate ligand, AA (**Figures 3A** and **B**), implying that these individual mutations can generically enhance BenM activity. However, the consensus mutations S, G, and H individually resulted in activity comparable to WT, indicating that these mutations were modulating BenM activity only in the context of other mutations (**Figure 3A**). The subtractive analysis on the CAVSGH consensus revealed more nuanced outcomes. CAVSGH had the desired biosensor behavior of high dynamic range for CCM (**Figure 3A**), including near-zero response in the absence of CCM and minimal sensitivity for the non-cognate ligand, AA (**Figure 3B**). However, when S, G, or H were individually removed, activity in the absence of CCM substantially increased, which was not the case when either A or V was removed (**Figure 3A**). This implies that S, G, and H enforce CCM-dependent activity and substantially improve dynamic range, whereas A and V generically increase BenM activity in the context of the other consensus mutations. Interestingly, removal of C from CAVSGH resulted in an inactive BenM. This combined with the observation that C alone increased generic BenM activity (**Figures 3A** and **B**) implies that C interacts with at least one other mutation in the consensus set in an epistatic manner. It is notable that the active biosensor, CCM-3 (**Figures 1C** and **2A**), does not contain C but does contain AVSGH, suggesting that the secondary mutations in CCM-3, namely A29T or D213G, compensate for the loss of C in the CAVSGH consensus. Although we did not characterize the contributions of all secondary mutations individually, we can conclude that the secondary mutations in CCM-1-6 contribute additional function to the biosensing properties of the consensus. This follows from the fact that CCM-7 is identical to the CAVSGH consensus (**Figure 1C**) and CCM-1-6, containing secondary mutations, all have greater dynamic range, higher response at saturating CCM concentrations, and larger operational range than CCM-7 (**Figure 2A**).

For AA biosensors, we comprehensively tested all 16 combinations of mutations found in the GTSC consensus (**Figure 3C** and **D**). We found that all individual mutations in the GTSC consensus are important for increased AA biosensor activity (**Figure 3D**). However, G and C showed activity increases in the presence of both CCM (**Figure 3C**) and AA (**Figure 3D**) whereas S and (to a lesser extent) T showed pronounced activity increases only in the presence of AA, suggesting that T and S are consensus mutations responsible for changing the ligand specificity of BenM to AA. When combined, GTSC exhibited high activity in the presence of AA but also appreciable activity in the absence of any ligand and in the presence of the non-cognate ligand, CCM (**Figure 3C**). This high background activity and/or low specificity was observed in all combinations of G, T, S, and C except the TS mutant, providing further evidence that T and S are primarily responsible for reprogramming BenM’s ligand specificity. Likewise, high background and low specificity was exaggerated in the GC double mutant, supporting the conclusion that G and C increase the activity of BenM in a generic manner. Overall, we observed that the functional contributions of mutations in the GTSC consensus could be qualitatively explained by an additive model wherein the activity of BenMs in the presence of AA largely increased with the number of consensus mutations and the background activity of BenMs was lowered by T and/or S, which also lowered activity in the presence of AA to a small extent in rare cases.

Secondary mutations found in the AA biosensors were important for further reducing the background activity of the GTSC consensus in the absence of ligand (**Figures 3E and F**). For example, G44D, T82I, I188N, S284P, and A294V, improved the dynamic range of GTSC by reducing background without substantially changing response in the presence of AA (**Figure 3F**). This suggests a model of evolution where there are a small number of large-effect mutations that increase the induced activity of BenM, and a more diverse number of small-effect background-lowering mutations accumulated throughout the mutational trajectories. This model is consistent with the higher stringency of positive selection steps versus negative selection steps during our evolution campaign. Taken together, these data reveal the subtleties of how OrthoRep-driven evolution achieved a diverse set of high-performance BenM biosensors that respond to CCM and AA and provide an additional collection of BenM variants with intermediate activities that may be useful in biosensor applications.

### Evolved BenM variants show broad operational range in presence of pathway enzymes producing CCM

To show that our evolved BenM variants are applicable in the metabolic engineering of CCM and AA production pathways *in vivo*, we tested the performance of our biosensors in the presence of an established biosynthetic pathway for the production of CCM in *S. cerevisiae* (16). The goal of this experiment was to demonstrate that our evolved biosensors 1) could properly report on the biosynthesis of CCM or AA and 2) possess an operational range that extends wide enough such that improvements in the biosynthesis pathway could be robustly detected in a high-throughput screen or selection. With these criteria in mind, we chose to test a subset of our CCM (CCM-3, CCM-4, and CCM-6) and AA (AA-3, AA-5, and AA-6) biosensors that exhibit high dynamic ranges, diverse operational ranges, and specificity for their cognate ligand (**Figures 1C** and **2**).

We genomically integrated the selected BenM variants (*ura3Δ0::REV1p-BenM_Variant-tTDH1*) into a strain containing 1) GFP driven by the hybrid CYC1p-BenO promoter and 2) a CCM biosynthesis pathway consisting of codon-optimized AroZ from *Podospora anserina* (*Pa*AroZ), CatA from *Candida albicans* (*Ca*CatA), and AroY from *Klebsiella pneumonia* (*Kp*AroY). As previously described, this particular combination of *Pa*AroZ, *Ca*CatA, and *Kp*AroY results in the efficient production of CCM in *S. cerevisiae* (16). While the addition of an enoate reductase (*ERBC* from *Bacillus coagulans*) can reduce CCM into AA, allowing us to also test our AA biosensors in the context of AA bioproduction, a multi-stage fermentation strategy utilizing a bioreactor in anaerobic conditions is required to produce active enoate reductase and consequent AA (24). For practical reasons, we decided to forgo direct production of AA, instead only supplying AA exogenously, which would still allow us to assess the operational range and specificity of AA biosensors in the presence of the upstream CCM pathway.

As shown in **Figure 4A**, CCM-3, CCM-4, and CCM-6 were all activated in the presence of the CCM biosynthetic pathway. They also all demonstrated the capacity for substantial further activation for the detection of improvements in CCM biosynthesis. For example, CCM-4 shows an additional ~6-fold activation in response to 7.04 mM exogenous CCM and an additional ~16.5-fold activation in response to 14.08 mM exogenous CCM. The benchmark TM biosensor is also capable of detecting endogenously biosynthesized CCM without saturating response, but only weakly senses the addition of exogenous CCM, while the benchmark MP17_D08 biosensor and WT BenM are both already saturated by the amount of endogenously biosynthesized CCM (**Figure 4A**). Compared to the benchmarks, our evolved CCM biosensors should therefore be more versatile in their application to the engineering of CCM biosynthetic pathways.

**Figure 4.**
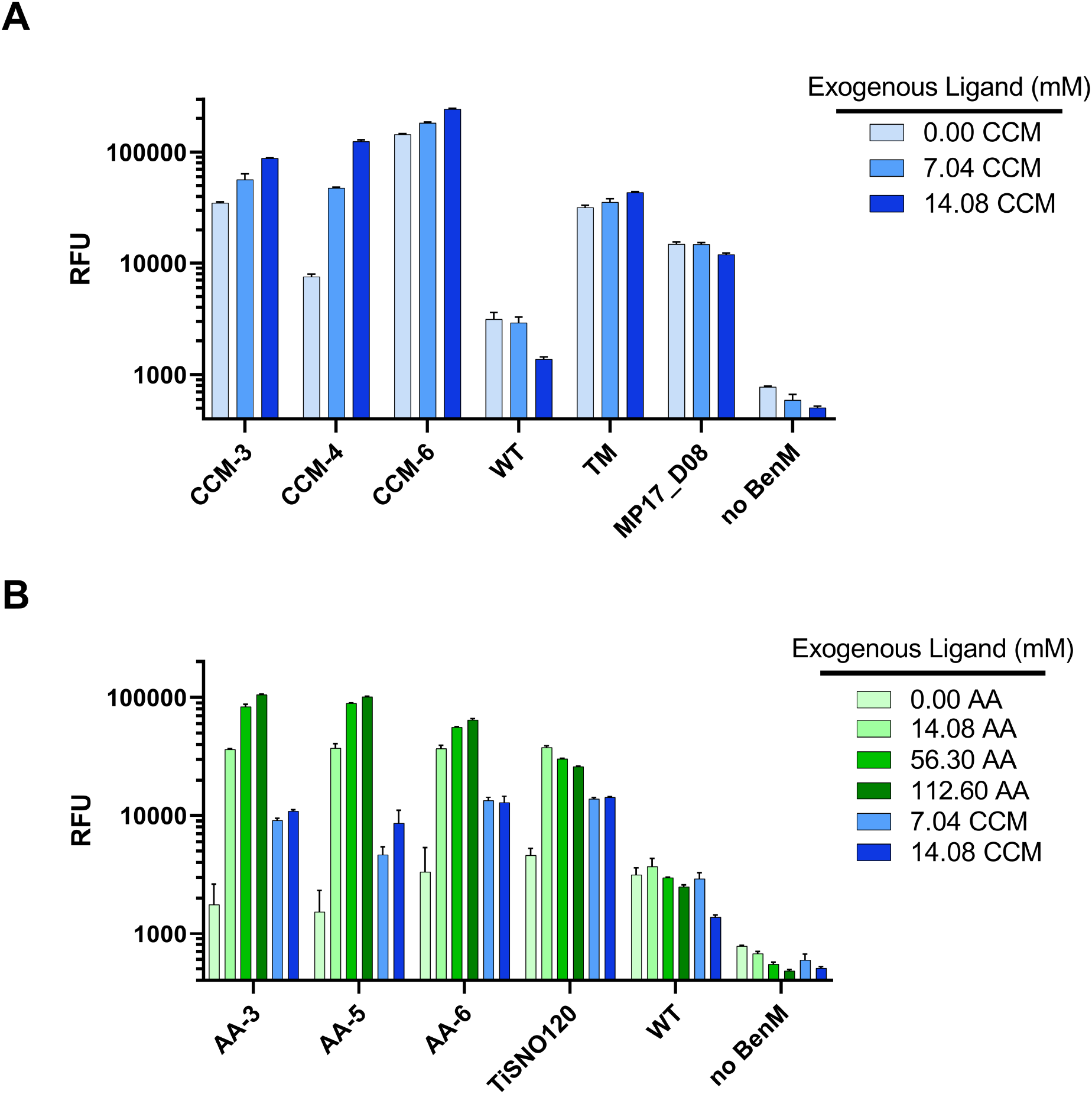
Performance of biosensors in strains producing endogenous CCM. (**A**) CCM biosensor response in a strain producing CCM with varying amounts of exogenous ligand added. (**B**) AA biosensor response in a strain producing CCM with varying amounts of exogenous ligands added. Exogenous ligand was added to culture media. Data were collected in biological triplicate, and the means ± one standard deviation (error bar) are shown.

A similar set of behaviors was found for our AA biosensors. As shown in **Figure 4B**, variants AA-3, AA-5, and AA-6 all exhibited significant further activation upon addition of AA in the presence of biosynthesized CCM, up to ~60-fold, ~65-fold, and ~19-fold for AA-3, AA-5, and AA-6, respectively. Additionally, even when 14.08 mM of exogenous CCM was used to supplement endogenously biosynthesized CCM, AA-3, AA-5, and AA-6 responded at levels well below their corresponding responses in the presence of 14.08 mM AA. This suggests that AA-3, AA-5, and AA-6 discriminate AA over high amounts of CCM, which should enable their application in engineering AA biosynthesis pathways that produce CCM as an intermediate. For comparison, the benchmark AA biosensor, TiSNO120, already exhibits saturation at AA concentrations below 14.08 mM. Overall, these data suggest that the BenM variants evolved in this study will be useful in the further optimization of CCM and AA production pathways in yeast.

## Discussion

We have shown here the successful generation of high performance *in vivo* aTF biosensors for CCM and AA using a directed evolution process where a parent biosensor is encoded on OrthoRep for continuous hypermutation *in vivo*. The biosensors we report have substantially improved dynamic range (up to ~180-fold in several cases), operational range, and ligand specificity compared to literature precedents, which should broadly expand the application space for CCM and AA biosensing, especially in the context of metabolic engineering. Indeed, we have demonstrated that our evolved biosensors successfully coupled to an existing CCM production pathway while retaining capacity for an additional ~16-fold activation with higher concentrations of CCM. Recently, Jensen *et al.* showed that OrthoRep could be used to drive the evolution of a biosynthetic enzyme in the CCM production pathway, using the MP17_08 BenM biosensor to guide selection for increased CCM titers (25). Since the BenM variants from this study have higher dynamic range than MP17_08 and are more effective at detecting high CCM and AA concentrations, they should be immediately applicable in the further evolution of CCM pathway productivity, as well as AA biosynthesis pathways that use an additional enzyme to convert CCM to AA. In the long run, we envision a general process where OrthoRep is first used to evolve aTF biosensors against desired small-molecules and then used to drive the evolution small-molecule production pathways with the biosensor to guide selection.

We attribute the high performance of our biosensors to the OrthoRep-driven evolution process through which they were generated. Specifically, OrthoRep effected the continuous hypermutation of BenM, allowing us to easily carry out many cycles of diversification and selection, because the process of evolution required only passaging of cells and FACS-based selection in the presence and absence of desired ligands. In this manner, our evolution campaigns were able to reach complex multi-mutant evolutionary outcomes that encompassed a variety of desired biosensor behaviors. The streamlined nature of biosensor evolution with OrthoRep is well matched to the demands of the broader biosensor engineering field. There are many small-molecules beyond CCM and AA for which effective aTF biosensors are lacking and for each distinct small-molecule, multiple concentration ranges need to be sensed. OrthoRep-driven evolution may prove uniquely scalable to match such breadth of target functions in the biosensor engineering space. Indeed, it may be possible to encode libraries of biosensor parents onto OrthoRep, such as those containing all natural LTTR effector-binding domains, in order to maximize the chance of finding initial biosensor activity against any small-molecule. Subsequent cycles of OrthoRep-driven evolution would automatically improve towards high dynamic range and hone towards desired operational ranges as demonstrated already with BenM. These directions will guide future efforts in our lab and others, with the ultimate goal of having effective *in vivo* biosensors for all small-molecules of interest.

## Materials and Methods

### DNA plasmid construction

Plasmids used in this study are listed in Table S1. *E. coli* strain Top10 was used for all the DNA cloning steps. All primers used in this study were purchased from IDT. All enzymes for PCR and cloning were obtained from NEB. All plasmids were cloned via Gibson assembly (26). Cloning of sgRNAs was performed as described by Ryan et al. (27).

### Yeast strains and Media/chemicals

All yeast strains used in this study are listed in Table S2. The auxotrophic selection marker used on p1 (*URA3*) was first fully deleted from the genome via CRISPR/Cas9 (27) before integrating genetic cassettes onto p1 or p1/p2 transfer through protoplast fusion. The auxotrophic selection marker for the nuclear plasmid containing TP-DNAP1-4-2 (*TRP1*) was also fully deleted, in this case after protoplast fusion. Spacer sequences were designed using Yeastriction v0.1 (28). Protoplast fusion was used to transfer OrthoRep into strain TrayBP-A7 and was performed as described before (20). Yeast strains were grown in standard media including YPD (10 g/L Bacto Yeast Extract; 20 g/L Bacto Peptone; 20 g/L Dextrose) and appropriate synthetic drop-out media (6.7 g/L Yeast Nitrogen Base w/o Amino Acids (US Biological); 2 g/L Drop-out Mix Synthetic minus the appropriate nutrients w/o Yeast Nitrogen Base (US Biological); 20 g/L Dextrose). CCM (Sigma #15992) and AA (Sigma #A26357) were dissolved directly into SC media. Afterwards, the pH was adjusted to 4.0 and the media filter-sterilized. CCM was found to be insoluble above 2 g/L (14.08 mM).

### Yeast Transformation

All transformations were performed via the high efficiency Gietz method (29). p1 integrations were performed as before (18, 20). For p1 integrations, 2-4 μg of plasmid was linearized by digestion with ScaI, which generated blunt ends containing homologous regions to p1. For *CEN6/ARS4* nuclear plasmid transformations, roughly 100-500 ng of plasmid was transformed. Combinatorial mutants were synthesized as gBlocks (IDT) and transformed directly into yeast. Transformants were selected on the appropriate selective solid SC media. Plates were grown at 30°C for 2 days for nuclear transformations and 4-5 days for p1 integrations.

### FACS-based evolution

Initial drift was performed by growing cells in 150 mL of SC-UW (pH 5.8) at 30 °C (250 r.p.m.) with daily 1:128 passages. After 21 passages (~150 generations), FACS-based screening began, consisting of alternating positive and negative sorts. After culture saturation (~20 hrs growth), cells were harvested and washed with a buffer containing HEPES-buffered saline (HBS; 20 mM HEPES pH 7.5, 150 mM NaCl) containing 0.2% maltose. After resuspending in buffer, FACS was performed with a Sony SH800 using a 70 μm Sony Sorting Chip. For positive sorting, cells were grown in SC-UW media (pH 4.0) containing 7.04 mM (rounds 1-6) or 14.08 mM (rounds 7-14) ligand (CCM or AA), whereas for negative sorting, cells were grown in SC-UW media (pH 4.0) without any ligand. Roughly 50,000,000 events and 20,000,000 events were measured for positive and negative sorting, respectively, for each evolution condition. Fluorescence was measured using a 488 nm laser and the top 0.5% fluorescence cells were recovered for positive sorts, while cells exhibiting background fluorescence were recovered for negative sorts. Cells were recovered in 40 mL of SC-UW (pH 4.0) until saturation (30 °C, 250 r.p.m.) and were subsequently diluted 1:100 into 40 mL of appropriate media, as described above, for the next round of selection.

### Flow cytometry

All measurements to characterize biosensor responses were taken by flow cytometry. Strains were streaked onto solid media and single colonies picked into 400 μL of SC or SC-UW media (pH 4.0). After 24 hrs growth (30 °C, 750 r.p.m.), cultures were diluted 1:100 into control (SC or SC-UW, pH 4.0) or induction media (SC or SC-UW + CCM/AA, pH 4.0). After 21 hrs growth (or 16 hrs for CCM-pathway containing strains), cells were diluted into 0.9% NaCl and measured on an Attune NxT Flow Cytometer (Life Technologies). Fluorescence of GFP was measured for 20,000 events, and the mean fluorescence for each population was determined. Fold-induction was calculated by dividing mean fluorescence of the induced condition (for a given concentration of ligand) by the mean fluorescence of the uninduced condition. To fit the titration curves, a non-linear least squares regression using a four-parameter model was performed in GraphPad PRISM. Each concentration measured corresponds to the mean of 2 or 3 biological replicates.

## Data Availability

The datasets analyzed are available from the corresponding author upon reasonable request.

## Supplementary Data

Supplementary Tables 1 (Table_S1.xls) and 2 (Table_S2.xls).

## Acknowledgements

We thank M. Jensen (Technical University of Denmark) and E. Jensen (Technical University of Denmark) for insightful discussions and the gift of the BenM gene, reporter strain MeLS0138, and CCM production strain ST4245. We thank M. Jensen for helpful edits and comments on our manuscript.

## Author Contributions

A.A.J. and C.C.L. were responsible for the conception of this study and experimental design. A.A.J. carried out all experiments. A.A.J. and C.C.L. analyzed data and wrote the manuscript.

